# Intercalation of the disulfide bond between the A2 and A4 loop of Cellobiohydrolase (Cel7A) of *Aspergillus fumigatus* enhances catalytic activity and thermostability

**DOI:** 10.1101/2023.07.13.548902

**Authors:** Subba Reddy Dodda, Musaddique Hossain, Sudipa Mondal, Shalini Das, Sneha Khator (Jain), Kaustav Aikat, Sudit S. Mukhopadhyay

## Abstract

Disulfide bond is important for maintaining the structural conformation and stability of the protein. Introducing new disulfide bond is a promising strategy for rational protein design. In this report, disulfide bond engineering has been applied to improve the stability of an industrially important enzyme, Glycoside Hydrolase family GH 7 cellobiohydrolase (GH7 CBHs) or Cel7A of *A fumigatus* origin. Disulfide by Design 2.0 (DbD2), an online tool, was used for the detection of the mutation sites and created four mutations (D276C-G279C; DSB1, D322C-G327C; DSB2, T416C-I432C; DSB3, G460C-S465C; DSB4) both inside and outside of the peripheral loops but, not in the catalytic region. The disulfide bond formed between the A2 and A4 loop of DSB3 showed higher thermostability (70% activity at 70^0^C), higher substrate affinity (K_m_= 0.081mM) and higher catalytic activity (K_cat_ =9.75 min^-1^; K_cat_/K_m_ = 120.37 mM min^-1^) compared to wild type *Af*Cel7A (50% activity at 70^0^C; K_m_= 0.128mM; K_cat_ = 4.833 min^-1^; K_cat_/K_m_ = 37.75 mM min ^-1^). The other three mutants with high B factor showed loss of thermostability and catalytic activity. Molecular dynamic simulations revealed that the mutation T416C-I432C makes the tunnel wider (DSB3:13.6 Å; Wt: 5.3 Å) at the product exit site; giving flexibility in the entrance region and mobility of the substrate. It may facilitate substrate entry into the catalytic tunnel and releases the product faster than the wild type. Whereas in other mutants, the tunnel is not prominent (DSB4), the exit is lost (DSB1), and the ligand binding site is absent (DSB2). This is the first report of the gain of function of both thermostability and enzyme activity of cellobiohydrolase Cel7A by disulfide bond engineering in the loop.

**Importance:** Cellulases are important for their role in the production of bioethanol, the cleanest renewable replacement of fossil fuels. Engineering of the cellulases is a chalange to increase their catalytic activity and thermostability for production of cheap ethanol. In this report we have introduced disulfide bond and successfully increased the both thermostabilty and catalytic activity of *Af*Cel7A.

## Introduction

The disulfide bond is a covalent link between the two sulfur (S-S) molecules in the cysteine residues of the proteins, which holds a significant role in making the structural conformation and function of the protein. Disulfide bond maintains the stability of the protein. However, the overall effect of the disulfide bond on the protein is complex. A general thermodynamic phenomenon is that the protein gains stability by reducing the conformational entropy of unfolded state. Introducing a new disulfide bond may increase the thermostability by disrupting the folding rate to the unfolding form of the protein (1). Engineered disulfide bonds have been widely used to improve protein stability, and several of them successfully create novel disulfides. But not all engineered disulfides produce an increase in stability. Even destabilising disulfides have also been reported. The factors that determine whether an engineered disulphide bond will increase or decrease the stability of a protein are not well characterised. Initial attempts were made on phage T4 lysozyme, subtilisin, dihydrofolate reductase, and lambda repressor for understanding the stabilising effect of engineered disulfides, and this information provides a general guideline for disulfide engineering technology (2–5). The main challenge in this strategy is to select the amino acid residue pair within the proper spatial distance, forming a covalent bond and giving stability.

A successful protein engineering prerequisite is the structural knowledge that retains the enzyme’s structure-functional relationship (6–8). The crystal structure of Cel7A of various fungal origins has been analysed (9). More or less, all the structures are similar, and it comprises the catalytic domain (CD), carbohydrate-binding module (CBM) and glycosylated linker. The catalytic domain is a tunnel-like structure comprising several peripheral loops (10). The crystal structure of the catalytic domain of *Af*Cel7A revealed that it consists of eight peripheral loops similar to *Tr*Cel7A(11). The flexibility and stability of these loops are crucial for substrate binding, hydrolysis, product expulsion, and processivity (12, 13). Various studies suggested the role of aromatic and polar residues of the CBM of Cel7A and the importance of glycosylation of the linker region (14). Recently Schiano-di-Cola et al., 2018 suggested that the loops are essential for enzyme activity by systematically deleting the flexible loops of the *Tr*Cel7A(10). Rational protein design by site-directed mutagenesis has been used to enhance the thermostability and catalytic activity of the Cel7A (6, 15–18). Cel7a has been engineered for substrate specificity (19), product inhibition (20), and ionic liquid tolerance (21). Insertion of disulfide bonds in an enzyme’s loop or flexible region can increase the enzyme’s thermostability (22–26). However, it may or may not hamper the catalytic activity.

There is a strong correlation between the thermostability and mobility of the mutated residue pairs used for disulfide bond engineering (27). This change in the mobility of amino acid residues can influence the catalytic activity depending on the mechanism of enzyme-substrate interactions, which may be positive or negative. The Ossowski IV. et al., 2003 introduced a new disulfide bridge (D241C/D249C) in the exo-loop of *Tr*Cel7a and achieved the activity on both amorphous and crystalline cellulose (8). The disulfide bridge mutation (G4C/M70C) near the N terminus and close to the entrance of the active site tunnel of *Ma* Cel7B showed improved thermostability ((28). Heterologous expression of all the s-s bridge mutations in or between the loops of the *Te* Cel7a showed thermostability except one (N54C/P191C), which offered improved activity (29). Here we have introduced disulfide bonds in the loop regions of *Af*Cel7A, not in the core catalytic domain.

The newly inserted disulfide bonds can influence the flexibility of the loops by which we can precisely corroborate how the flexibility of the loops plays a pivotal role in the catalytic activity of the enzyme *Af*Cel7A. DbD2 (disulfide by designing 2), a computational tool based on B factor determination is used to predict the possible sites and these sites are replaced by cysteine to create disulfide bonds. The B-factor strongly correlates with the stability of the mutated residues and the enzyme. The high B-factor indicates surface exposed residues (27, 30). *Af*Cel7A contains nine disulfide bridges, and eight of them are conserved (31, 32). Among different suggested mutants we selected four mutants (D276C-G279C; DSB1, D322C-G327C; DSB2, T416C-I432C; DSB3, G460C-S465C; DSB4) based on the B factor values. The mutations are mainly in the peripheral flexible loop regions surrounding the catalytic domain but not in proximity to the old disulfide bridge. The recombinant mutant enzymes expressed in *Pichia pastoris* under an inducible promoter (AOX1) and further purified for biochemical characterisation. The mutant T416C-I432C/DSB3 formed a disulfide bridge between the A2 and A4 loop and showed higher thermostability and catalytic activity than wild-type. Our approach of disulfide bond engineering between the A2 and A4 loop of *Af*Cel7A is the first report, which enhances both the thermostability and catalytic activity. However, the other three mutants having relatively high B factor lost the thermostability and catalytic activity compared to wild-type *Af*Cel7A. The molecular dynamic simulation study elucidated the structure-function relationship of the enzymes and their mutated variants.

## Results

### Screening of potential Disulfide bond mutants

Out of nine native disulfide bonds of *Af*Cel7A eight (164C-430C, 87C-93C, 202C-235C, 45C-51C, 76C-97C, 198C-236C, 287C-363C, 256C-282C, 264C-269C) are conserved with other fungal species. The DBD2, the web-based tool, identified 35 possible mutants for creating disulfide bonds (Supplementary Table. 1). Only four mutations (D276C-G279C/DSB1, D322C-G327C/DSB2, T416C-I432C/DSB3, G460C-S465C/DSB4) were selected based on the B-Factor score. Disulfide bonds form between 3-16 amino acids and do not interfere with the native disulfide bonds. All the mutants have mostly positive Chi3 (χ3 torsion angle), indicating the proper dihedral angle of the disulfide bond. All the designed disulfide bonds are 2.7nm to 6nm to the active site residues. No abrupt structural changes were observed in the mutated and other locations of the enzymes. The circular dichroism data suggest an almost unaltered secondary structure for mutant DSB3 and wild type (Table S4). T416C-I432C/DSB3 is forming the disulfide bond between A2 and A4 loops. D276C-G279C/DSB1 is forming a disulfide bond within the B3 loop. Whereas the G460C-S465C/DSB4 and D322C-G327C/DSB2, the mutations (point) are outside the loop regions of the *Af*Cel7A (Fig 1a & 1d). The model structure also infers the position of mutation sites, loops, selected residues, substrate, and enzyme interaction sites of the *Af*Cel7A (Fig. 1b & 1c).

**Figure 1.**
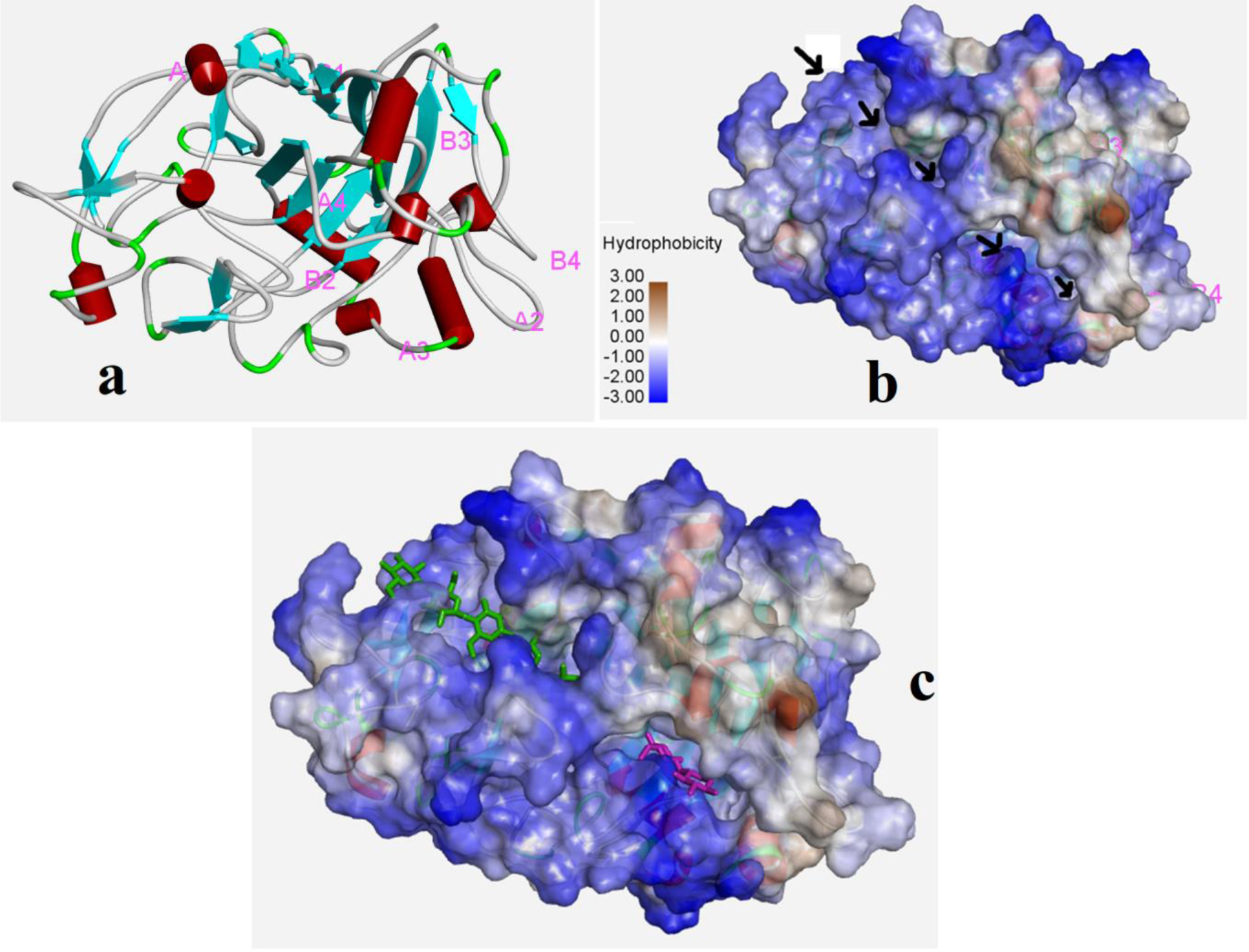
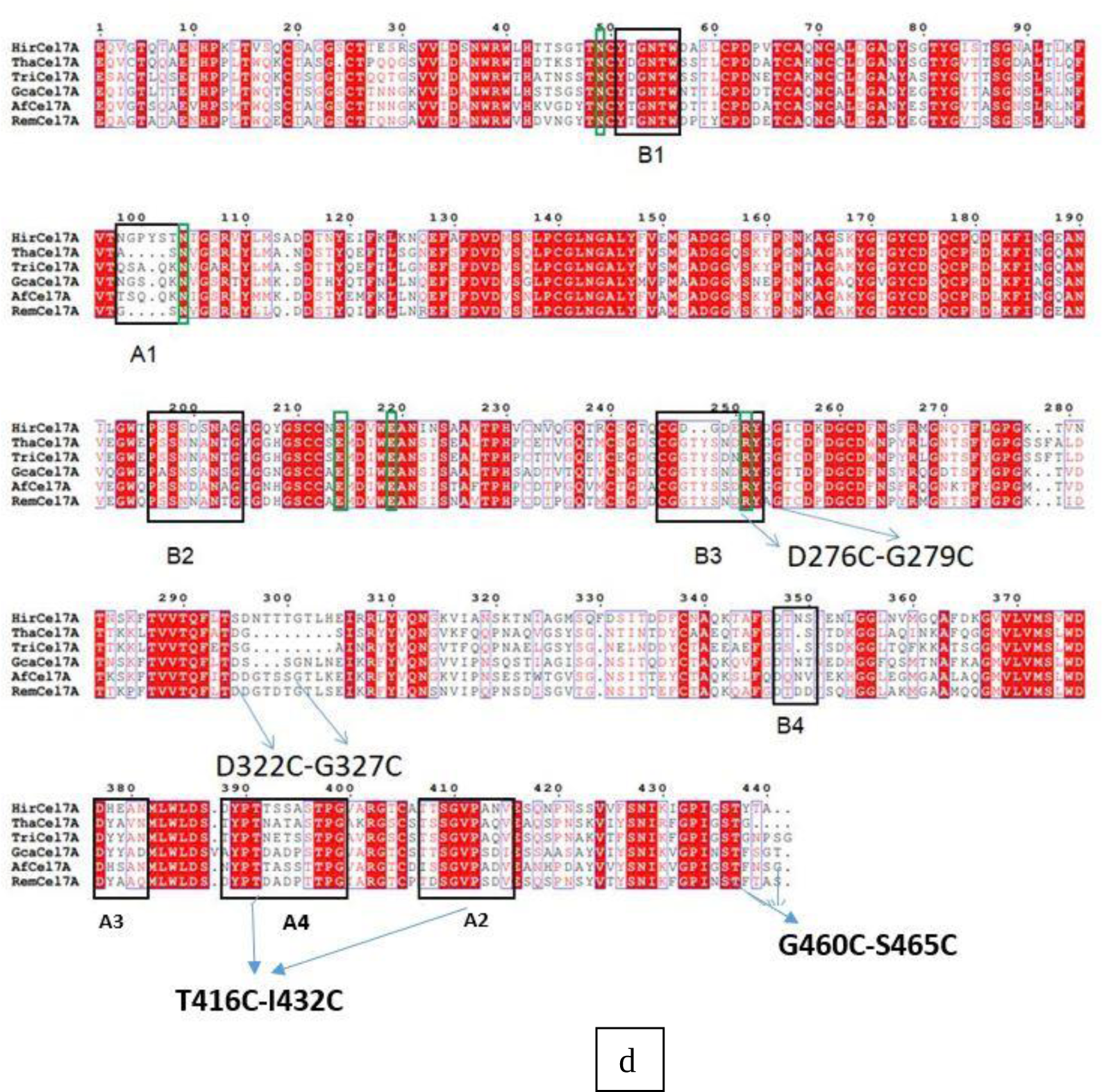
Representation of point mutation and the disulfide bond positions. (a) Modelled structure of wild-type protein, (b) the surface of the wild-type protein showing the tunnel, (c) beta cellohexaose and beta cellobiose inside the tunnel of wild type. **(d)** Multiple sequence alignment and indication mutation site (in every positioning, 26 amino acids should add because the alignment is of catalytic domains only).

### Mutation creation, purification, and disulfide bond confirmation of *Af*Cel7A

*Af*Cel7A was cloned in the pPICZαA vector under AOX1 (alcoholic oxidase) promoter. The targeted mutations were developed in a wild-type construct using the site-directed mutagenesis method (Supplementary Table 2) (33). The wild-type and the mutants were expressed in *Pichia pastoris* GS115 cells with 0.5% methanol induction. The enzymes were purified by Ni-NTA affinity chromatography, followed by the confirmation of a single band in SDS-page and Western blot (Fig. 2A and 2B). Enzymatic activity was determined using different substrates like pNPL (4-nitrophenyl β-d-lactopyranoside), Avicel, and biomass.

**Figure 2.**
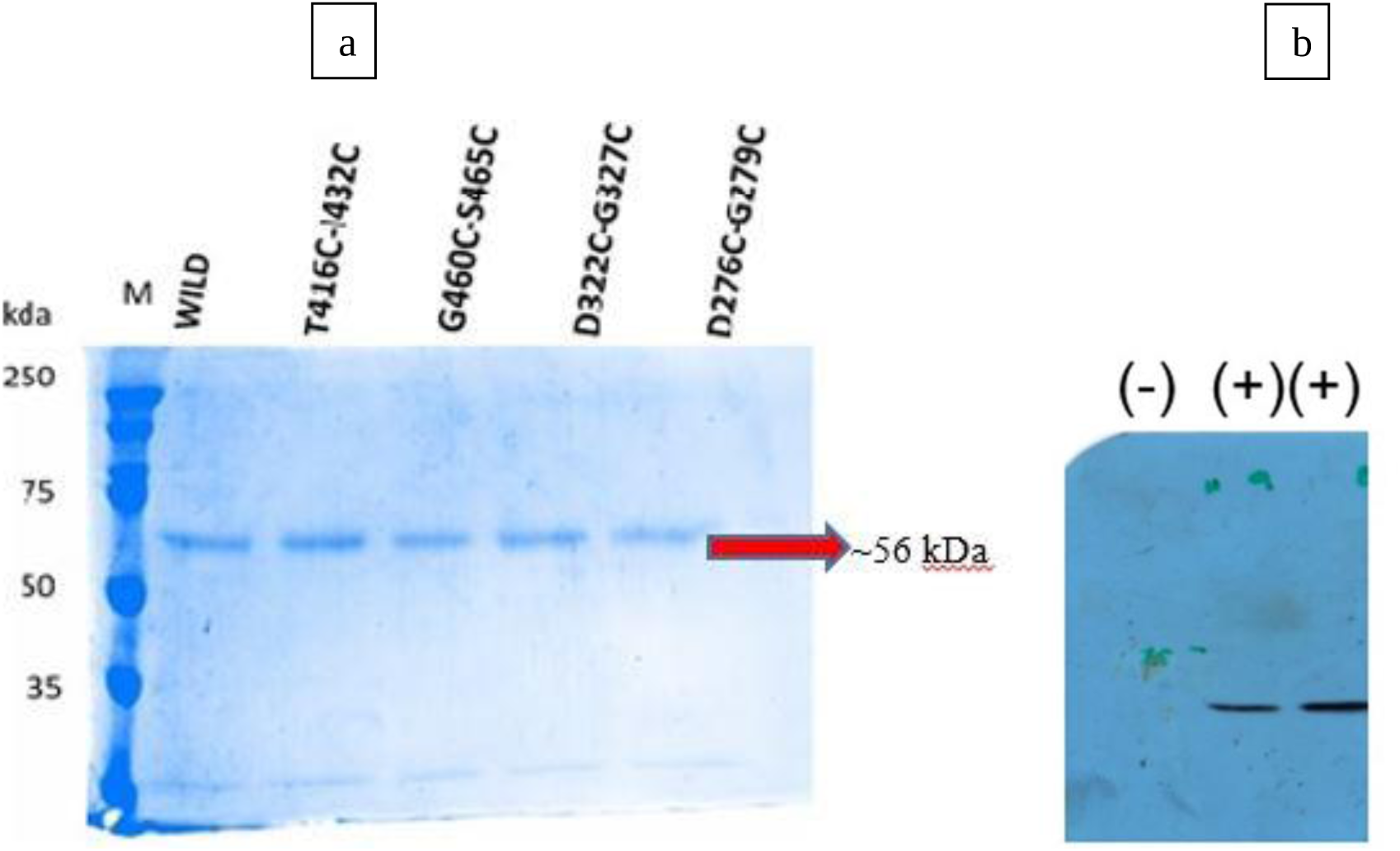
Expression and purification *Af*Cel7A and its mutants. **(a)** SDS-PAGE (12%), purified *Af*Cel7A, and mutants by Ni-NTA chromatography showing the band at approximately 56kDa. **(b)** Western blot of the AfCel7A and mutant T416C-I432C mutation and negative control (only *Pichia* secretory proteins)

In proteins, thiol groups (mercaptans or sulfhydryl) are in cysteine residues, and disulfide bonds are formed between the thiol groups. Therefore, detecting the thiol group suggests the amount of sulfhydryl in the protein. Thiol or sulfhydryl group was determined from the equal concentration of wild-type and mutant proteins using the well-established DNTB (Ellman’s reagent) method. The average concentrations of the thiol group are almost similar (∼300μM) in all mutant enzymes, which are relatively higher than the wild type (Fig. 3). These results confirmed the formation of new cysteine residues, which ultimately form the disulfide bond in the mutants, but not in the wild type.

**Figure 3.**
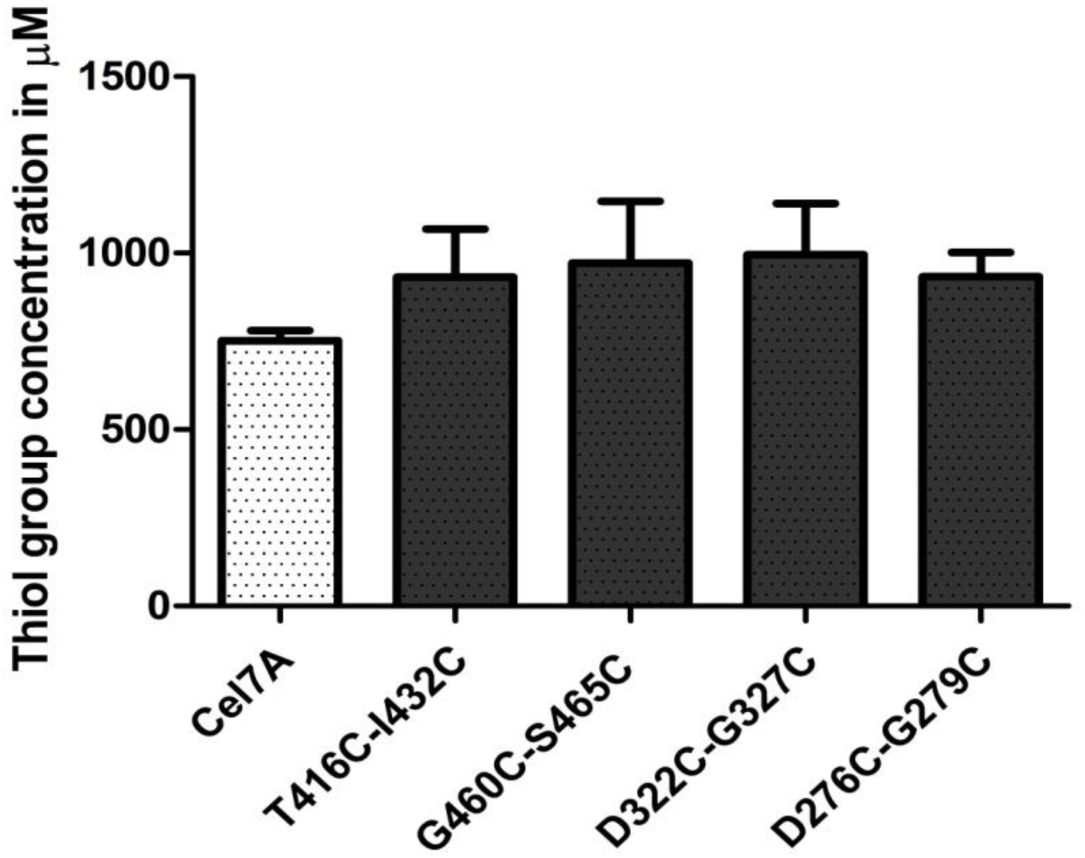
Free thiol groups or sulfhydryl group detection of *Af*Cel7A and its mutants measured by DTNB method.

### Thermostability of the mutant enzymes

The thermostability of the mutant enzymes (DSB1, DSB2, DSB3, and DSB4) was determined by identifying the relative activity at different temperatures and compared with wild-type *Af*Cel7A. The purified enzymes pre-incubated at different temperatures (30ᵒC to 70ᵒC) for 30 minutes, and the enzymatic activity was carried out in optimal conditions (pH 5 and 50ᵒC)(11). Interestingly the mutant enzyme T416C-1432C (DSB3) retained nearly 70% activity while pre-incubated at 70^0^C temperature for 30 minutes. The wild-type *Af*Cel7A showed almost 50% relative activity when pre-incubated at 70^0^C temperature for 30 minutes. However, the other three mutants (G460C-S465C, D276C-G279C, and D322C-G327C) failed to retain their thermostability compared to the wild type (Fig. 4).

**Figure 4.**
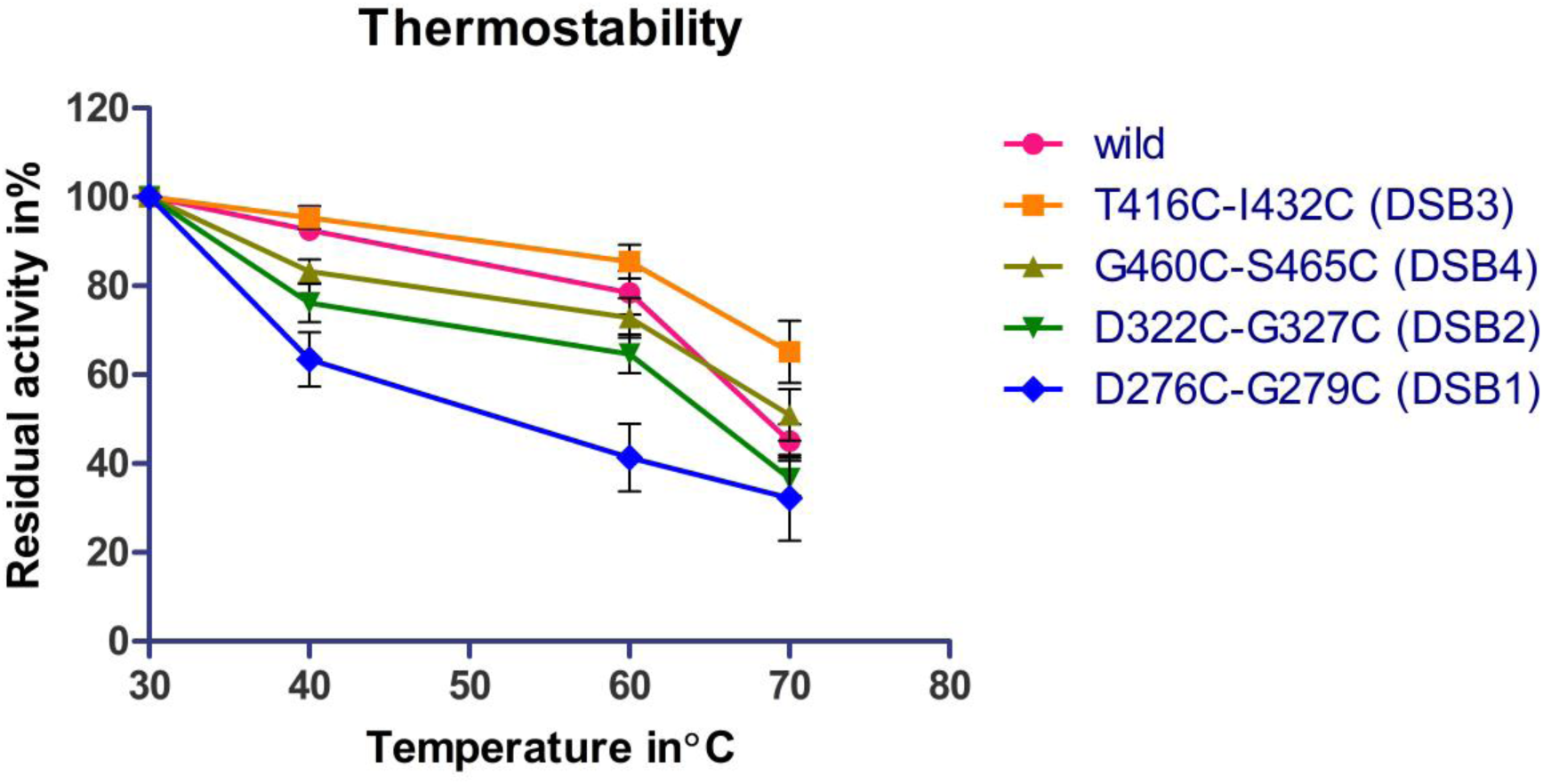
Thermostability of the *Af*Cel7A and its mutants. Redisulalacitvity of the wild Cel7A and its variants at varying temperatures in ᵒC using substrate (1.6mM pNPL(pH 5.0) for 10 mins)

### Enzyme kinetics and product inhibition study of *Af*Cel7A and its mutants

Here we used pNPL as substrate and performed the enzyme kinetics using various substrate concentrations. Simultaneously we conducted the enzyme inhibition study utilising the cellobiose as an inhibitor (product inhibitor). The Lineweaver-Burk plot was used to determine the enzyme kinetics parameters. All the parameters were compared between wild-type and mutants *Af*Cel7A (Table 1) (Supplementary Figure 5). The mutant T416C-1432C/DSB3 showed a higher catalytic rate constant (K_cat_ 9.75 min^-1^) compared to the wild type (4.833 min^-1^), and the Michaelis constant (K_m_) is (0.081mM) lower than wild type (0.128mM). These results confirmed the increase of substrate affinity and catalytic activity of the mutant T416C-1432C/DSB3 compared to the wild type. Other variants (DSB1, DSB2, and DSB4) showed the loss of substrate affinity (higher K_m_) and catalytic activity compared to the wild type (Table 1). Enzyme activity was also compared with the natural substrate (Fig 5) and biomass (Fig S3). When crystalline cellulose Avicel was used, DSB3 showed significantly higher activity than the wild type. The activity increased with the concentration. In 8 mg/ml concentration, DSB3 showed almost double activity compared to the wild type. Whereas the mutant DSB4 lost its activity compared to the wild type. It confirmed our results with the synthetic substrate pNPL. Similar results were also observed when we used natural biomass for hydrolysis. DSB3 showed significantly higher activity than the wild type, and DSB4 showed substantially lower activity (Fig S3).

**Figure 5.**
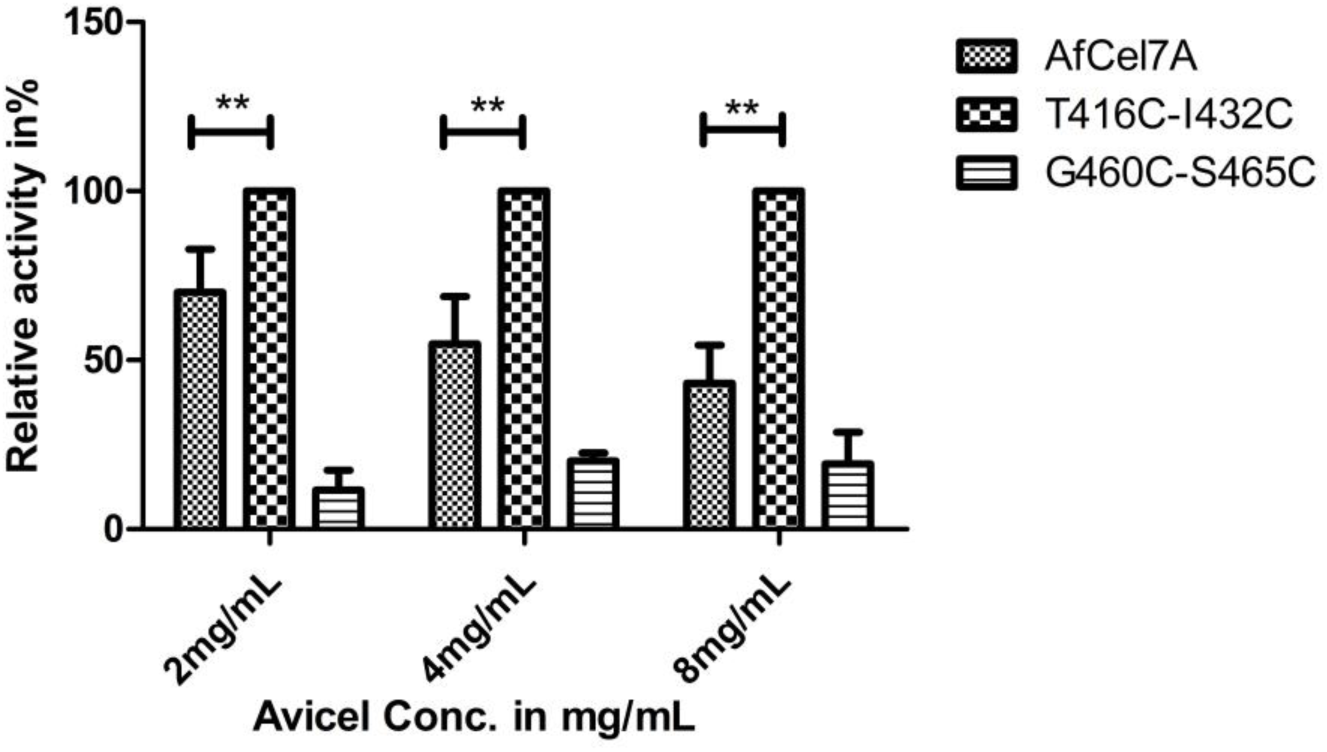
Relative activity plot: Relative activity of *Af*Cel7A, the mutant T416C-I432C and the mutant G460C-S465C with different Avicel concentration

**Table 1:** Kinetic characterisation of *Af*Cel7A and its mutants on pNP-Lac

| Enzymes | $K_m(\text{mM})$ | $K_{cat} (\text{min}^{-1})$ | $K_{cat}/k_m(\text{mM min}^{-1})$ |
| --- | --- | --- | --- |
| Wild cel7A | $0.128 \pm 0.001$ | $4.833 \pm 0.02$ | $37.75 \pm 0.2$ |
| T416C-I432C | $0.081 \pm 0.009$ | $9.75 \pm 0.05$ | $120.37 \pm 5$ |
| G460C-S465C | $0.75 \pm 0.03$ | $3.88 \pm 0.01$ | $5.17 \pm 0.3$ |
| D322C-G327C | $1.96 \pm 0.2$ | $3.36 \pm 0.1$ | $1.71 \pm 0.05$ |
| D276C-G279C | $1.24 \pm 0.5$ | $3.13 \pm 0.2$ | $2.52 \pm 0.25$ |

Cellobiose (disaccharide), the released product of *Af*Cel7A, inhibits enzyme activity. In this experiment, we have compared the product inhibition of mutants along with wild-type enzymes. We observed that the *Af*Cel7A and mutant T416C-1432C/DSB3 showed a nearly competitive inhibition pattern on the LB plot. However, the mutant G460C-S465C/DSB4 showed a way of non-competitive inhibition (Fig. S5). The inhibition constant value (K_i_) for wild-type *Af*Cel7A is 30μM. For the mutant DSB3, the K_i_ is 10μM. In contrast, the K_i_ for DSB4 is 50μM. Interestingly, the mutant DSB3 gained catalytic activity but lost product inhibition property. However, the mutant DSB4 lost the catalytic activity but gained the product inhibition property.

### Molecular Dynamic simulations

Computational structural studies provide a reliable description of the enzyme’s three-dimensional structure, enzyme-substrate interaction, and mobility details of the enzyme-substrate complex. MD simulation studies are also considered a complement to experimental studies (34, 35). Here we have performed the homology modelling followed by the MD simulation study (up to 100ns) to get an explanation of the biochemical behaviour of the *Af*Cel7A and its mutants. The MD simulations of the *Af*Cel7A-cellobiose complex and their mutants D276C-G279C (DSB1), D322C-G327C (DSB2), T416C-I432C (DSB3), and G460C-S465C (DSB4) were performed at 300K (Fig 10). The RMSD value of wild-type protein is 33.04Å, whereas the RMSD values are 31.85, 33.20, 33.28, and 32.9 for DSB1, DSB2, DSB3, and DSB4, respectively (Fig 10). We performed MD simulation (100ns) of all the wild types and the mutated enzymes by developing structure using homology modelling. A transparent tunnel is visible in the cavity of the wild-type enzyme (Fig. 1). This tunnel is comparable with the reported crystal structure of Cellobiohydrolase I (PDB id 7CEL) (Fig.S1). This tunnel is also present in the modelled structure of mutated protein DSB3 (Fig. 6), but it looks slightly wider and might be more flexible than the wild type. As per the wet lab experiments, the catalytic activity and the substrate affinity of the DSB3 are higher than that of the wild-type. Here the tunnel is slightly wider in the binding place of cellobiose. The product exit tunnel shows a cavity width of almost 13.6 Å, whereas the wild type showed a width of 5.3Å (Fig. 12a and 12b). In the case of DSB4, this ligand binding tunnel is present but not prominent like the wild type and DSB3 (Fig7 & 12C). A small tunnel is present in the mutated protein DSB1, which has an entrance for the substrate. Still, there is no opening for the product release (Fig. 8). For the mutated protein DSB2, this ligand binding tunnel is absent (Fig. 9). The radius of gyration (Rg), defined as the distribution of atoms of a protein around its axis. The R_g_ values of T416C-I432C (DSB3) and G460C-S465C (DSB4) are 32.7Å and 32.9 Å respectively, higher than the R_g_ value of the wild type, which is 30.8Å. But the R_g_ value of D276C-G279C (DSB1;15 Å) and D322C-G327C (DSB2;28.82 Å) are less than that of the wild type. RMSF plot of all the mutated proteins and wild type has been given in Fig. 10. It is observed that the overall fluctuations of the amino acids in mutated proteins are more compared to wild type. Fluctuations are higher in the case of DSB2 and DSB3. The structure of all the proteins is optimised and minimised by running calculations involving molecular dynamic simulation. Energy vs time graphs for the wild type, DSB3, and DSB4 have been plotted in Figure 11. RMSD and RMSF values (20ns) were also compared at different temperatures (Fig S4). Overall all the data suggests that the mutant DSB3 (T416C-I432C) is more stable than others.

**Figure 6.**
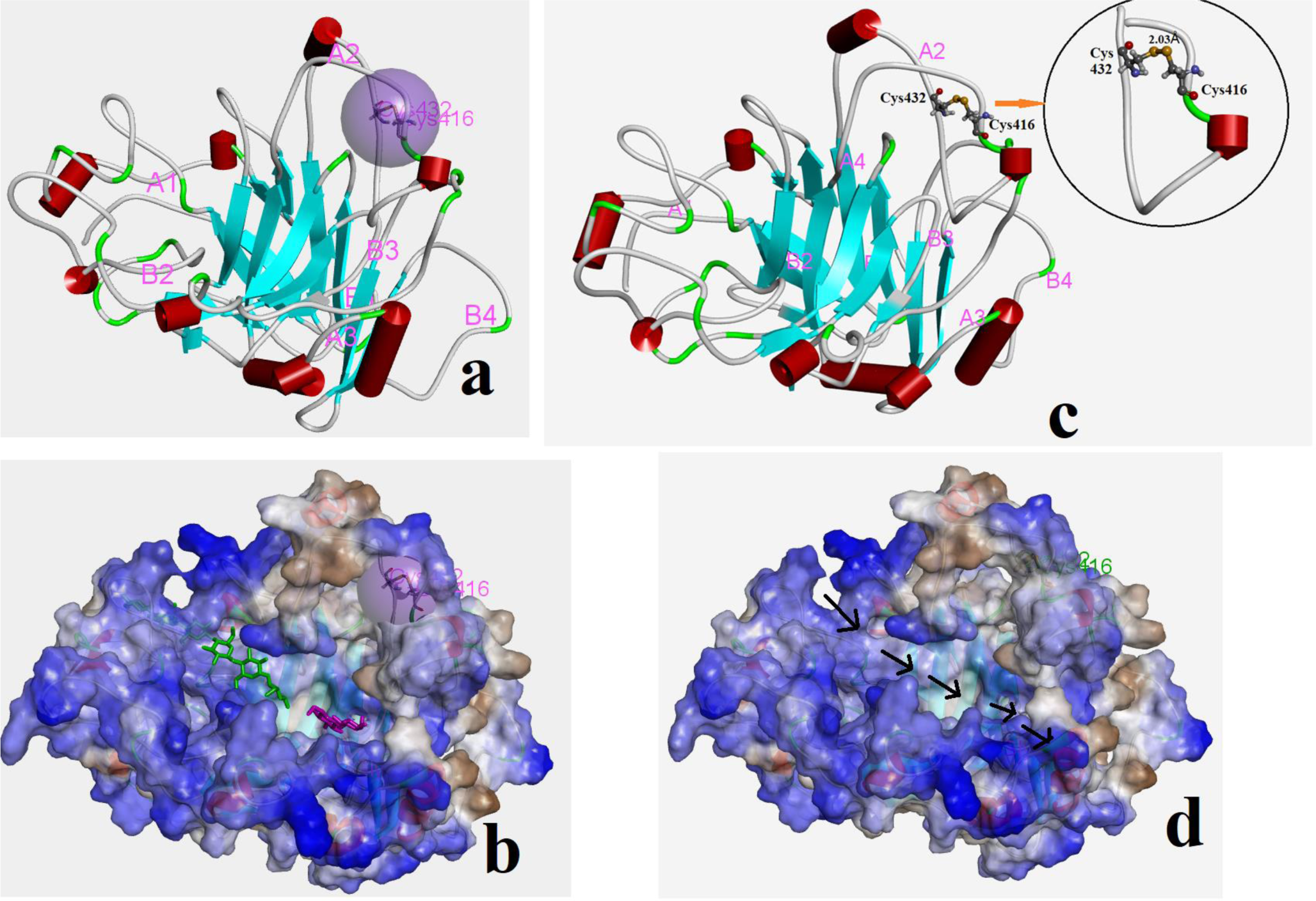
: (a)Modeled structure of DSB3, (b) Modeled structure of DSB3 focusing the S-S bond, (c) surface of the mutated protein and beta cellhexaose and beta cellobiose inside the tunnel. (d) tunnel in the mutated protein DSB3.

**Figure 7.**
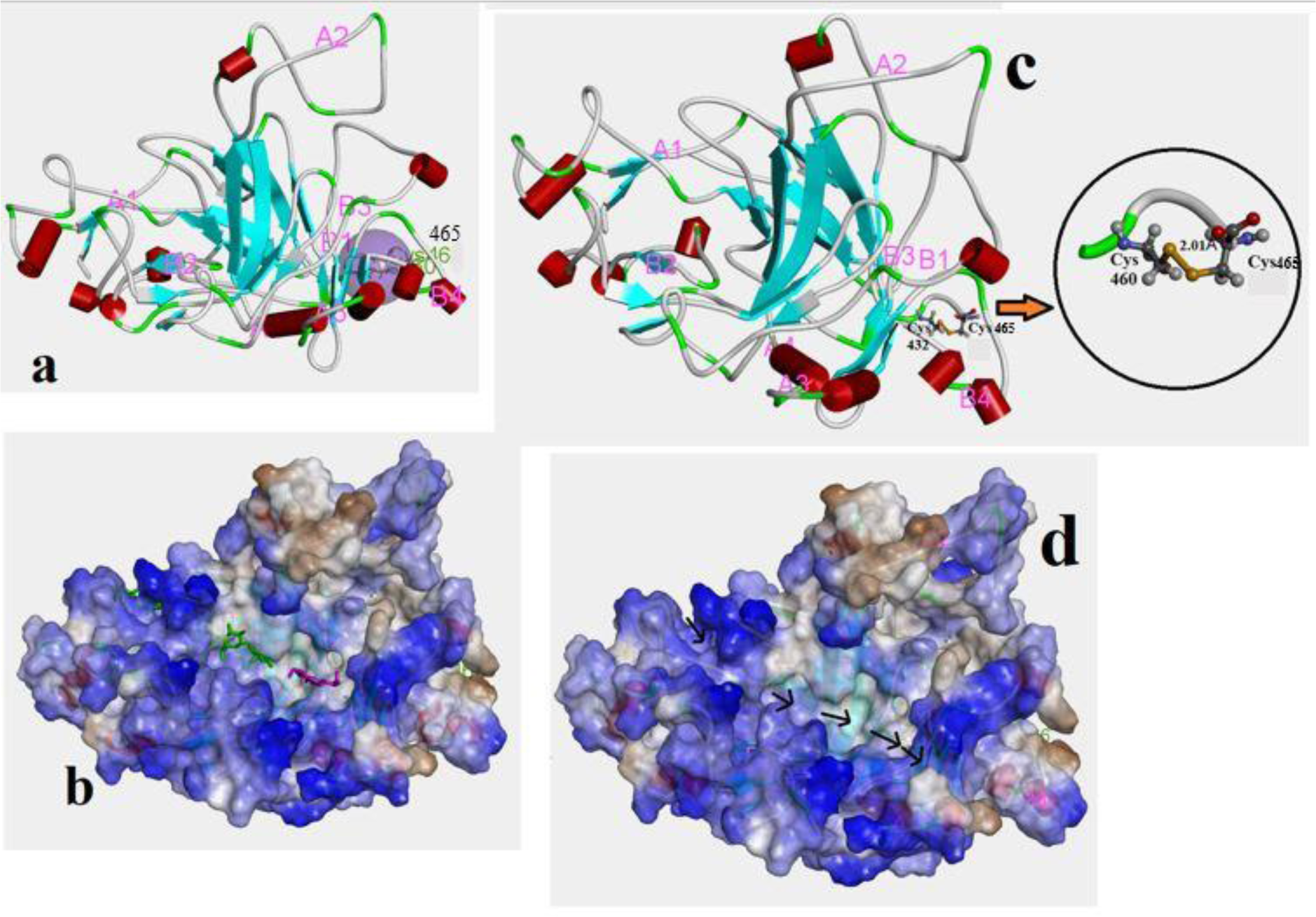
(a)Modeled structure of DSB4, (b) Modeled structure of DSB4 focusing the S-S bond, (c) surface of the mutated protein and beta cellhexaose and beta cellobiose inside the tunnel.(d) tunnel in the mutated protein DSB4.

**Figure 8:**
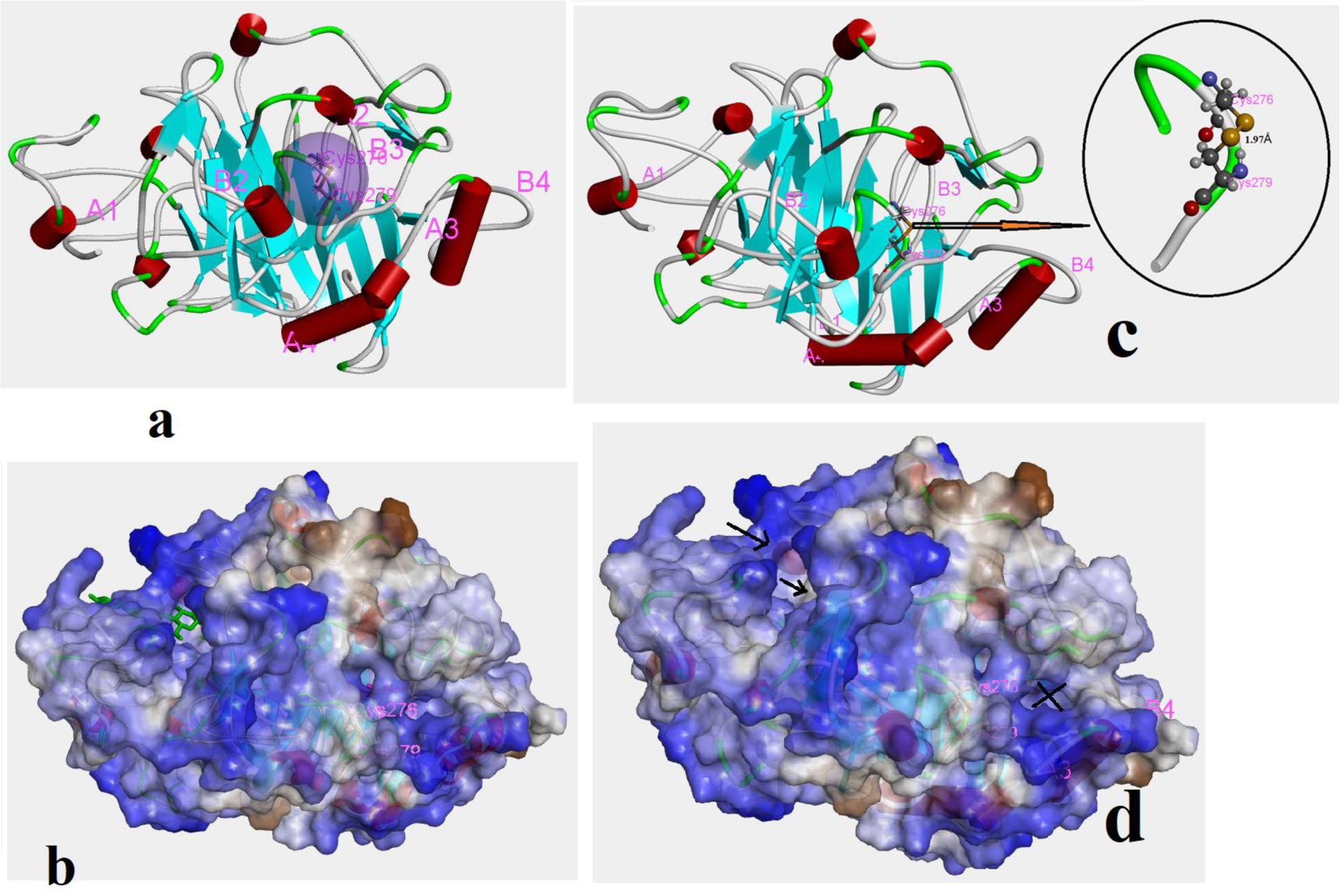
(a)Modeled structure of DSB1, (b) Modeled structure of DSB1 focusing the S-S bond, (c) surface of the mutated protein and beta cellhexaose and beta cellobiose inside the tunnel.(d) tunnel in the mutated protein DSB1.

**Figure 9:**
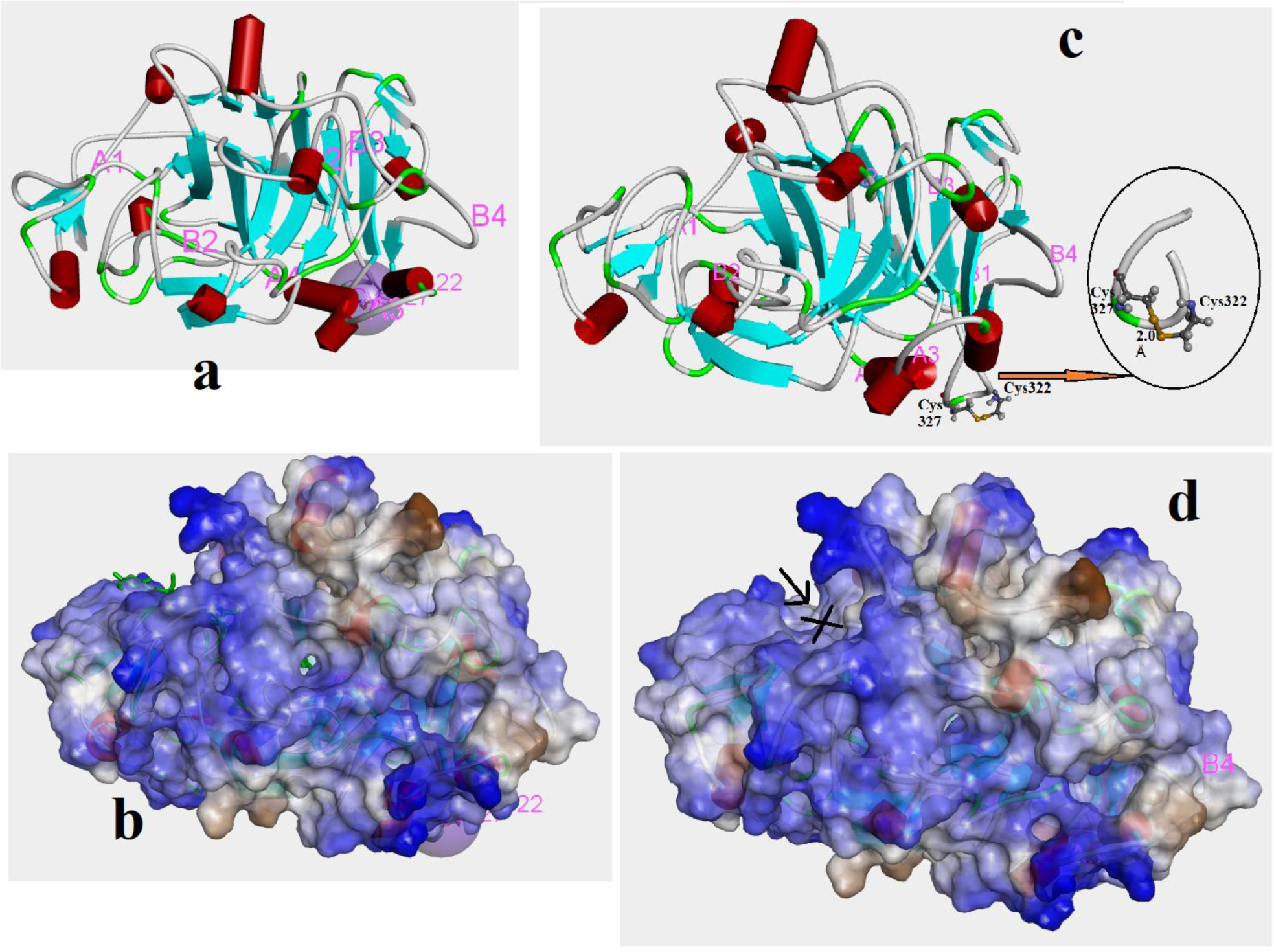
(a) Modeled structure of DSB2, (b) Modeled structure of DSB2 focusing the S-S bond, (c) surface of the mutated protein (d) tunnel in the mutated protein DSB3 where beta cellhexaose and beta cellobiose cannot go inside the tunnel.

**Figure 10:**
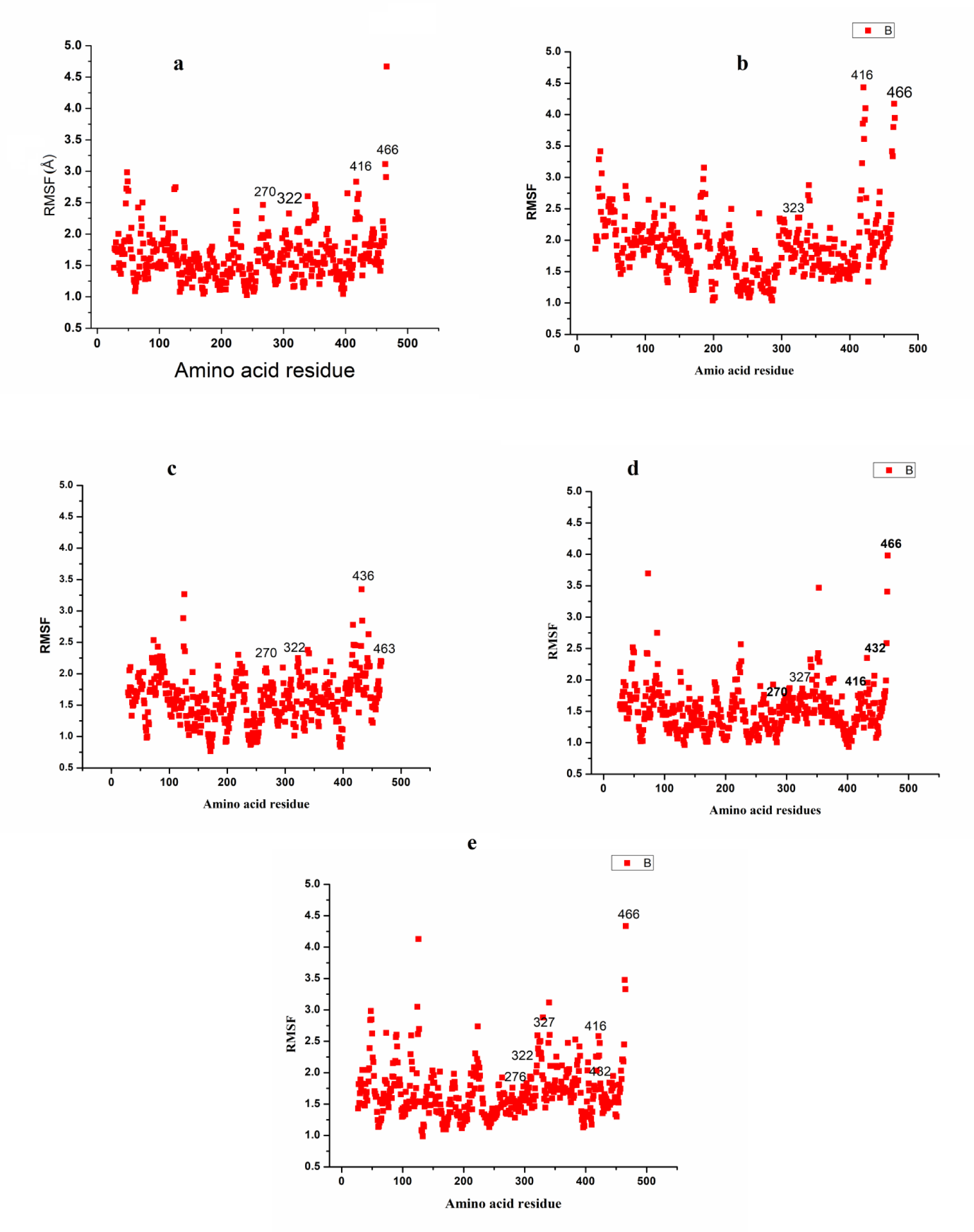
Plot of RMSF vs amino acids residues (a) wild type, (b) DSB3, (c) DSB4, (d) DSB1, (e) DSB2.

**Figure 11:**
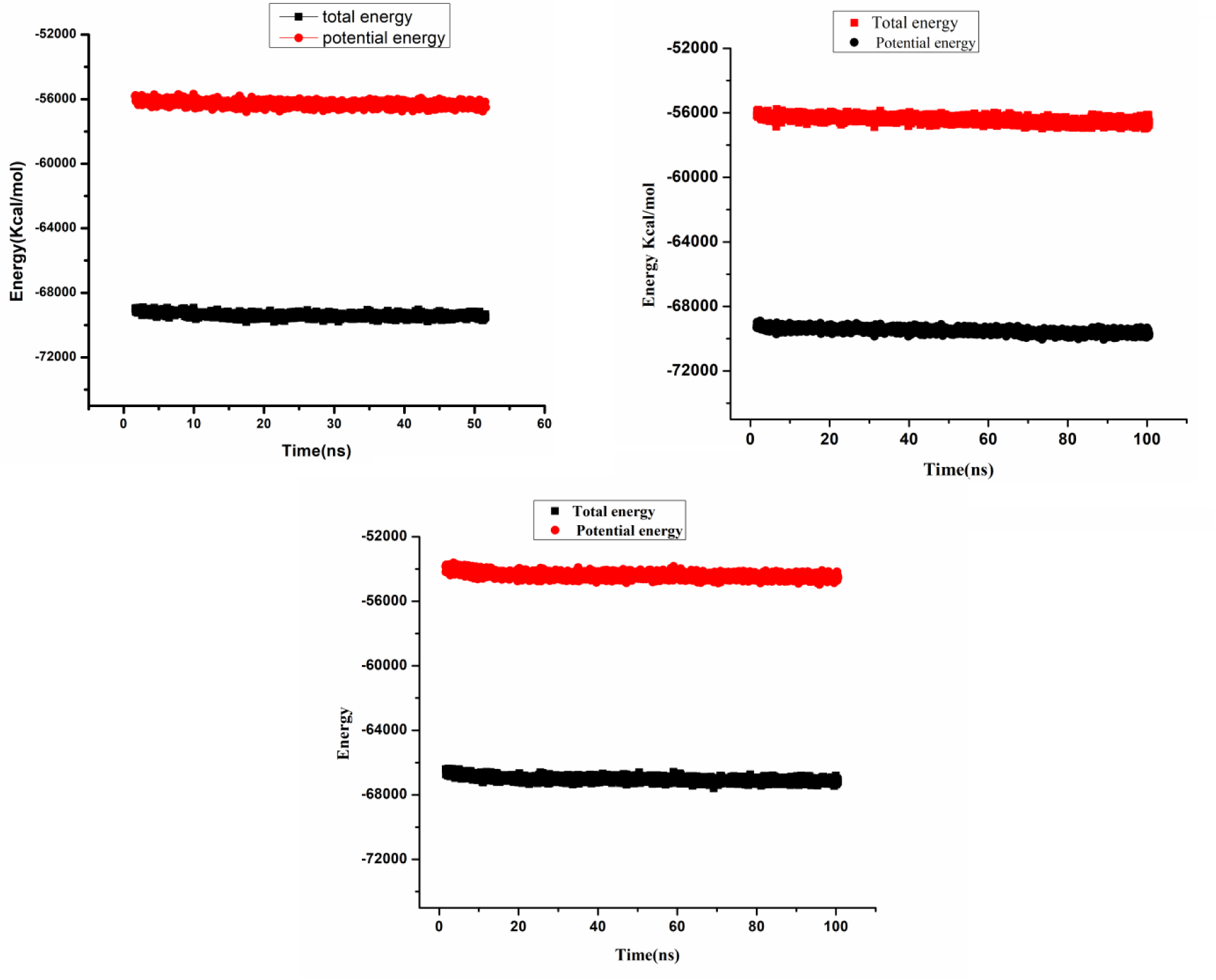
Plot of Energy vs time of *Af*Cel7A and mutants. (a) wild type, (b) DSB3, (c) DSB4.

## Discussion

Stability and catalytic activity are the most important characteristics of an industrial enzyme. Disulfide engineering is a potential approach for designing novel disulfide bonds into the target protein to enhance stability. But the factors determine the disulfide bond-mediated stabilising effect of the protein are poorly characterised. We have screened the mutation sites based on the B-factor (DbD2), which is crucial in maintaining the mutant protein’s stability and the mutant residue’s three-dimensional geometrical positioning. The high B-factor dealt with the higher stability of the mutant protein and the surface exposed residues (27). Here, we observed that the mutations having high B-factor (>60) (D276C-G279C/DSB1, D322C-G327C/DSB2, and G460C-S465C/DSB4) (Supplementary Table 1) showed lower thermostability and on the other hand, the mutant T416C-I432C/DSB3 with comparatively lower B-factor (<50) showed higher thermostability compared to wild type (Fig 4). Similarly, when enzyme activity was measured by determining the enzyme kinetics with synthetic substrates like pNPL and enzymatic activity for natural substrates like Avicel or natural biomass, we observed that the mutations having high B factor (DSB1, DSB2, and DSB4) lost their activity. Still, a mutant with the lower B factor (DSB3) showed higher activity than the wild type (Table 1, Fig 5 and Supple Fig 3). On the contrary, when the cellobiose inhibition was determined, the DSB4 (high B factor) showed higher K_i_ (50µM) than the wild type (30µM). Whereas DSB3 (lower B factor) showed a lower K_i_ value compared to the wild type (10µM) (Supplementary Fig.5). The k_i_ values suggest that the enzyme activity of the mutant T416C-I432C/DSB3 is inhibited by a lower concentration of the cellobiose compared to wild type. But the activity of the G460C-S465C/DSB4 is inhibited by higher concentration of cellobiose compared to the wild type. So, DSB3 has become more sensitive to cellobiose, and DSB4 has become resistant to cellobiose compared to the wild type. These results revealed a tradeoff mechanism between the catalytic activity and product inhibition in *Af*Cel7A. The probable molecular mechanism behind the gain of thermostability and catalytic activity can be explained through the MD simulation study. The reported crystal structure clearly shows that the catalytic domain in cellobiohydrolase is like a tunnel. The tunnel has an entrance at one end and an exit at the other end. The substrate enters through one end, releasing the product through the other end (Fig.S1). Crystal structure represents the cellohexose and cellobiose binding sites at the two places inside the catalytic tunnel. Two clear binding sites are also observed in the cavity of the wild-type protein; the bigger cavity is for the cellohexose, and the smaller one is for the cellobiose (Fig.S2). The mutation T416C-1432C (DSB3) makes the tunnel wider; specifically, the exit has become three-fold wider than the wild type (Fig 12b). This gives better flexibility or mobility of the entrance region and may facilitate the entry of substrate into the catalytic tunnel and releases the product faster than the wild type. An immediate insight into the structure of DSB3 shows that the A4 loop has moved upward from the ligand-binding site due to the S-S bond formation between Cys416 and Cys432. This displacement of the A4 loop in DSB3 may cause the wideness of the tunnel. Here the flexibility in the entrance region of the tunnel is comparable with the wild type. However, the binding position of beta cellobiose is wider in DSB3. Also, it contains more hydrophobic residues (Fig. 6b). However, the scenario is quite different in the mutant G460C-S465C (DSB4). Here S-S bond formed between Cys460 and Cys465, so loops are in different positions. A detailed insight into the structure of DSB4 displays that the B3 loop is near the binding site of cellobiose. Still, in the case of wild-type and DSB3, the B3 loop is away from the binding site, and it contains more hydrophobic residues (Fig.1b, 6b). Hence, the lower activity of DSB4 can be explained by the disposition of the B3 loop. In the case of DSB1, there is no proper exit in the catalytic tunnel and the product (cellobiose) release is hindered. This is due to the formation of the S-S bond in Cys276 and Cys279. A close vision of the structure depicts that the S-S bond may hinder the formation of the exit of the tunnel. We have observed that the radius of gyration of DSB1 is 28.15 Å, which is less than that of the wild type (30.8Å). This indicates that DSB1 is more compact in structure and may partially abolish the catalytic tunnel (Fig 8). For DSB2, the S-S bond has formed between Cys322 and Cys327, and the loops have different rearranged positions that cause the absence of the tunnel (Fig 9). Here the position of A1 and B2 loops are in such a position that the entrance of the tunnel is almost closed (Fig. 9d). The radius of gyration of DSB2 is 28.82 Å which also represents that this mutated protein is more compact than the wild type and may be a reason for the lack of binding space for cellohexaose and cellobiose. The plot of energy vs time states that the total and potential energy of the wild type and DSB3 are almost (Fig. 11a, b) same but DSB4 is energetically unstable compared to the wild type and DSB3. So, residues in the catalytic loop are crucial in controlling the catalytic activity, substrate affinity, and product inhibition. This study also suggests that the amino acid residues’ position and the substrate-enzyme interaction mode are the most critical parameters for designing disulfide bond engineering. Finally, we can conclude that this is the first evidence that we have successfully employed the disulfide bond engineering method for improving the thermostability and catalytic activity of the GH7 family of Cellobiohydrolase of *Aspergillus fumigatus* origin.

**Figure 12:**
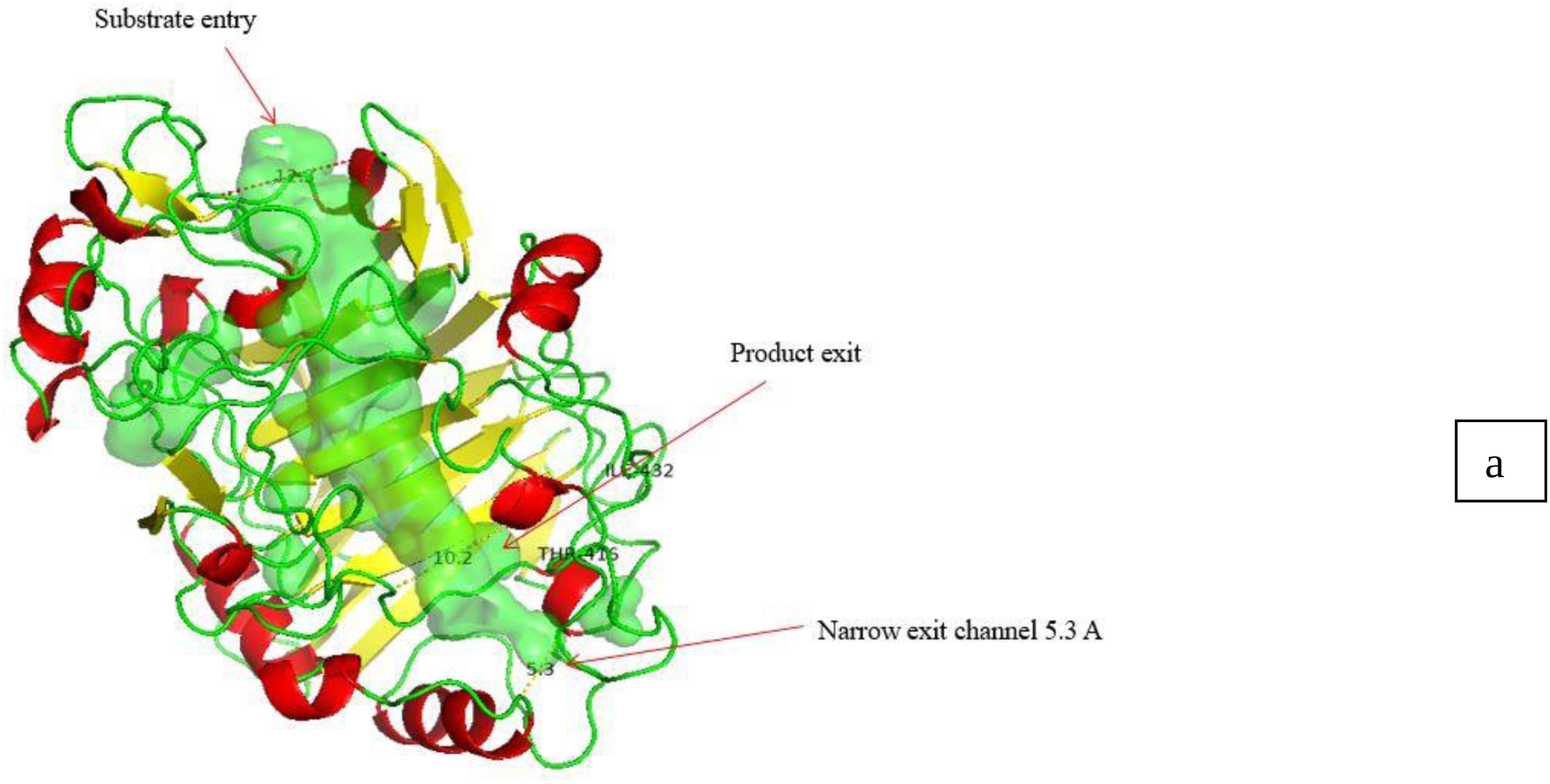
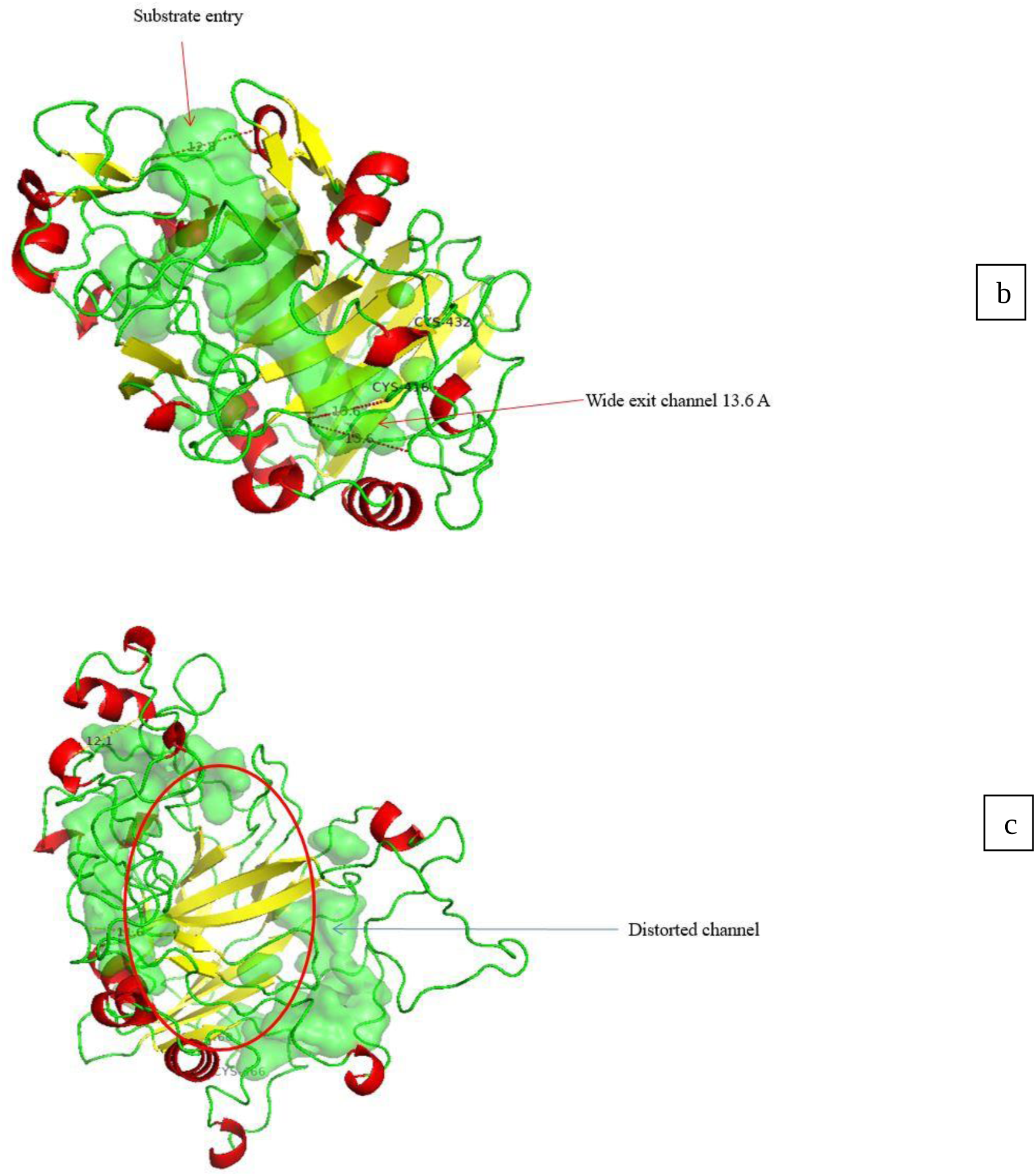
The substrate entry channel and product exit region; (a) *Af*Cel7A the channel marked in green. (b) DSB3: the wider product exit region ∼13Å. (c) DSB4: Distorted channel

## Experimental procedures

### Designing of Disulfide bond mutants

Disulfide by design (DbD) is a program to identify the residue pairs likely to create the disulfide bridges (27). The 3D structure of the *Af*Cel7A (PDB Id: 4V1Z) is given as an input. The mutants were created by using primer-based site-directed mutagenesis (36).

### Cloning and expression of Cel7A and its variants

*AfCel7A* gene was amplified by PCR from the genomic DNA of *A fumigatus* NITDGPKA3. The primers were designed based on the sequence available in NCBI (gi|71025133). The primers of wild type and mutants are given in supplementary table 2. The PCR product was cloned into the *EcoR*1 and *Not*1 sites of the vector pPICZαA (Invitrogen) and transformed into *E. coli* DH5α cells. The positive colonies were confirmed by colony PCR, restriction digestion, and sequencing. The linearised pPICZα-Cel7A was transformed into *P. pastoris* GS115 (Easy Select kit, Invitrogen). The positive colonies were selected based on colony PCR and were expressed for three days with 0.5% methanol. The culture supernatant was filtered and precipitated by Ammonium Sulfate precipitation (50%). Protein was purified by IMAC (Immobilised affinity chromatography) chromatography (37). The concentration of protein was estimated by the Bradford method using BSA as a standard. The protein was separated on 12% SDS-PAGE gel and stained with Coomassie brilliant blue R250. Western blot further confirmed the purified protein with an anti-His monoclonal antibody (Abcam, USA).

### Enzyme Activity Assays

Enzyme activity was determined with Avicel (increasing concentration starting with 2mg/mL to 8mg/mL), and the reaction was carried out with 50mM sodium-phosphate buffer ph-5.0. The pNPL (1.33mM *p*-NP-β-d-lactopyranosid) reactions were carried out with 25mM citrate buffer pH-5.0. The reaction solution contained 100µl of a diluted enzyme, 500µl of Avicel, and 400µl of a suitable buffer. The reaction solution was kept at 50°C for 1 hour with continuous stirring. The reducing sugar estimation was done by the Dinitro salicylic acid assay (DNS) method(38), and glucose was used as a standard. The amount of enzyme that produces 1µM of glucose per minute under standard assay conditions was considered one unit. The optimal pH and temperature were determined under standard conditions using pNPL as a substrate. Alkaline pre-treated raw rice straw was used as a substrate for biomass hydrolysis assay. The 50 mM sodium-phosphate buffer ph-5.0 was used as the standard reaction buffer. The biomass and Avicel dissolve in the same buffer. The mutant enzyme T416C-I432C activity was taken as 100% (highest), and the activity of other enzymes was shown relatively with the mutant one. In each reaction, enzyme blank and substrate blank are set as control.

### Thermostability assay

As the industry prefers, we have determined indirect activity-based thermostability, not direct protein stability. All the purified enzymes are incubated at different temperatures starting from 30^0^C to 70^0^C for 30 minutes. The substrate *p*-NP-β-d-lactopyranoside (pNPL) (1.33mM) was incubated with the enzyme in 25mM citrate buffer at optimum reaction conditions (pH 5 and 50ᵒC) for 30 minutes. The amount of pNP released was measured and compared with the enzymes at the optimum reaction temperature. The activities at different temperatures were expressed as relative activity (%), considering the untreated enzyme activity as 100%.

### Enzyme kinetics and cellobiose inhibition

The enzyme kinetics was studied using *p*-NP-β-d-lactopyranoside (pNPL) as a substrate. The substrate was added with increasing concentration (0, 0.13, 0.26, 0.66, 1.3, 2.6, 5.3, 6.6 mM) and was incubated with appropriately diluted enzyme (100µl) at 50 °C for 30 min. The reaction was terminated by adding 100µl of sodium carbonate (500 mM). The product inhibition was studied in the presence of cellobiose (20 µM and 80µM). All reactions were performed in triplicate, and kinetics data were analysed through the Line Weaver Burk plot. The product released from the pNPL was measured using a p-nitrophenol standard curve. The linear regression fitting was performed using Graph Pad Prism to determine Vmax, Km, and Kcat. Graph Pad Prism was also used for statistical analyses. The inhibition constant (K_i_) was calculated by the equation Vo=Vmax[S]/(α*K*m+[S]) where α=1+[I]/KI where I is the concentration of inhibitor.

### Disulfide bond identification *in vitro*

The presence of a disulfide bond was examined by Ellman’s reagent 5, 5’-Dithiobis (2-nitrobenzoic acid) (DTNB) method(39). This method quantitatively determines the number of free sulfhydryl groups in protein structure. Protein (mg/ml) was dissolved in 0.1 M sodium phosphate buffer (pH 8.0) and aliquoted into two. One aliquot was treated with DTT (30mM), while the same amount of 100mM sodium phosphate buffer (pH 8.0) was added to another one. The samples were incubated at 25^0^C for 2 hours. Then the samples were dialysed against 0.1 M sodium phosphate buffer (pH 8.0) overnight to remove the DTT and salts. Protein samples were concentrated and measured by the Bradford method. 100μl of DTNB was added to 400ul of reduced and non-reduced protein samples, and mixtures were incubated at 25^0^C for 5 minutes. Absorbance was measured at a 412nm wavelength to monitor aromatic thiolate release. The number of thiol groups was determined by coefficient value E412nm TNB2– = 1.415 × 104 cm^-1^M^-1^. A sample without treating DTT was considered a negative control.

### Circular dichroism spectroscopy

CD spectra were recorded at room temperature on a Jasco J815 Spectropolarimeter controlled with J-800 software. The spectra were obtained at a protein concentration of 0.1mg/ml in a 40mM phosphate buffer using a 1 mm path length. The data was recorded in triplicate using wavelength from 260 to 190 nm range at a scan rate of 50nm/min using a 2.0-sec response and a sensitivity standard by the software. The data pitch was 1nm with a 2nm bandwidth. The data measures circular dichroism along Y-axis in deg and along the X-axis wavelength in nm. High tension voltage (HT) is also measured along a different Y-axis. CD data were analysed on BEST SEL’s online tool (http://bestsel.elte.hu/).

### Sequence alignment and Molecular Dynamics simulation

Sequences of all the Cellobiohydrolase were retrieved from the PDB database (https://www.rcsb.org/), and alignment was carried out by the Clustal Omega(40). The graphical image of sequence alignment was generated by the ESPript 3.0 server(41). Discovery Studio Visualizer 2022 produced superimposed structures(42)^-^(43). To get the 3D structure of the mutated proteins, homology modelling was done using the macromolecule module of Discovery Studio software package 2022. Many crystal structures of different cellobiohydrolases were available in the protein data bank. Still, the structures with PDB id 4V20, 3PFZ, 4ZZV, and 4ZZP were taken as templates since they showed the highest sequence identity with minimum E value with good resolution. Calculations involving MD simulations were performed to get the mutated proteins’ energy-minimised structure by the Discovery Studio software simulation module. Before the MD simulation, the proteins were solvated with a water molecule in an explicit periodic boundary in TIP3P water molecules. The solvated proteins were minimised first, followed by the heating for 120ps and equilibration at NVT ensemble for 1000ns. After equilibration, the protein structures were simulated. Wild type protein was simulated for 50ns but mutated protein T416C-I432C (DSB3) and G460C-S465C(DSB4) for 100ns at 300 K with a time step integration for 2fs in the NVT ensemble. MD simulation was performed by applying Charmm force field. The structure analyses like RMSD, RMSF, R_g_ were performed using Analyze trajectory toolbar. The pictures are generated by PyMol(43).

## Author contributions

SRD and SSM conceptualised the whole research; SRD, MH and SD performed the experiments, and SM did the theoretical calculation and data interpretation. SRD, SM, SSM and MH wrote the manuscript.

## Acknowledgement

SRD is thankful to the DST-INSPIRE, DST and DBT, Govt. of India for providing fellowship. The authors are grateful to the DST-FIST grant of the Department of Biotechnology, NIT Durgapur. The authors also thank Bose Institute and Dr Ajit Bikram Dutta for providing the necessary facilities. Intellectual property India patent (application number 202131050361 dated 02-11-2021)

